# mDia2 formin selectively interacts with catenins and not E-cadherin to regulate Adherens Junction formation

**DOI:** 10.1101/721530

**Authors:** Yuqi Zhang, Krista M. Pettee, Kathryn N. Becker, Kathryn M. Eisenmann

## Abstract

**Background:** Epithelial ovarian cancer (EOC) cells disseminate within the peritoneal cavity, in part, via the peritoneal fluid as single cells, clusters, or spheroids. Initial single cell egress from a tumor can involve disruption of cell-cell adhesions as cells are shed from the primary tumor into the peritoneum. In epithelial cells, Adherens Junctions (AJs) are characterized by homotypic linkage of E-cadherins on the plasma membranes of adjacent cells. AJs are anchored to the intracellular actin cytoskeletal network through a complex involving E-cadherin, p120 catenin, β-catenin, and αE-catenin. However, the specific players involved in the interaction between the junctional E-cadherin complex and the underlying F-actin network remains unclear. Recent evidence indicates that mammalian Diaphanous-related (mDia) formins plays a key role in epithelial cell AJ formation and maintenance through generation of linear actin filaments. Binding of αE-catenin to linear F-actin inhibits association of the branched-actin nucleator Arp2/3, while favoring linear F-actin bundling. We previously demonstrated that loss of mDia2 was associated with invasive single cell egress from EOC spheroids through disruption of junctional F-actin.

**Results:** In the current study, we now show that mDia2 has a role at adherens junctions (AJs) in EOC OVCA429 cells and human embryonic kidney (HEK) 293 cells through its association with αE-catenin and β-catenin. mDia2 depletion in EOC cells leads to reduction in actin polymerization and disruption of cell-cell junctions with decreased interaction between β-catenin and E-cadherin.

**Conclusions:** Our results support a necessary role for mDia2 in AJ stability in EOC cell monolayers and indicate a critical role for mDia formins in regulating EOC AJs during invasive transitions.

## Background

Ovarian cancer is the deadliest gynecological malignancy, with 14,070 women estimated to die from the disease in the United States in 2018, according to the American Cancer Society. There are at least four types of ovarian cancer classified by their cell type of origin, with epithelial ovarian cancer (EOC) representing 90% of diagnosed cases [1]. Most patients are diagnosed in the late stages, when peritoneal dissemination has already occurred [2], and ∼75% of patients develop relapsing disease after undergoing current standard of care treatment with cytoreductive surgery and adjuvant platinum/taxane duplet combinations [3].

While hematogenous metastasis does occur, EOC’s primary route of dissemination is transcoelomic [4, 5]. Cancer cells are shed from the primary tumor as single cells or clusters into the peritoneal cavity where they are carried with the accumulated peritoneal fluid (ascites) to secondary sites such as the liver and diaphragm (as reviewed in [4]). Since cancer cell detachment from the primary tumor initiates metastasis, it is important to understand the molecular mechanisms involved in this process.

A type of cell-cell junction present in epithelial cells is the Adherens Junction (AJ), which typically consists of E-cadherin and p120, α-, and β-catenin [6-8]. AJ formation is calcium-dependent. In the presence of calcium, the extracellular domains of E-cadherin change conformation to allow for homotypic linkages between cells whose adjacent membranes are ∼20 nm apart [6, 9]. AJ stability is dependent on AJ protein anchorage to the underlying cortical F-actin cytoskeleton and continuous F-actin polymerization [10, 11]. While the classical AJ indicates direct linkage between the E-cadherin/β-catenin complex to F-actin via α-catenin, binding to β-catenin may reduce the affinity of α-catenin for F-actin [8, 12]. A recent alternative model proposes that α-catenin cycles between two intracellular pools-one junctional, where it associates with the cadherin complex- and another cytosolic or perijunctional that associates with the underlying F-actin [13]. The two α-catenin pools may allow for α-catenin to act as a molecular switch-turning off Arp2/3-mediated branched actin-polymerization to promote linear actin polymerization [13, 14]. The exact role of cortical F-actin in the adherens junction and its regulation by various catenins and actin-associated proteins is still a subject undergoing active investigation.

Various actin-associating and/or polymerizing proteins are known to affect AJ stability and are regulated by the catenins. Among these are various members of the formin family of proteins, which act as downstream effectors of Rho-GTPases. Upon binding of a Rho-GTPase to the GTPase-binding domain (GBD) of the formin, the formin is released from its autoinhibited state, promoting non-branched F-actin polymerization, and, in some cases, bundling [15]. Formin family members such as formin-1, Diaphanous-related formin 1 (mDia1), and Fmnl3 localize to and/or strengthen the AJ by increasing E-cadherin junctional localization while decreasing its mobility within the plasma membrane [11, 16, 17]. Formin localization to AJs was regulated through Rho-signaling and post-translational modifications such as phosphorylation and myristoylation [17-21]. Formins may also be mechanosensitive, responding to external forces. mDia1-mediated F-actin polymerization increased upon pN force application to actin filaments [22].

Both formin inhibition and activation were associated with increased cellular invasion. For example, mDia formins were required for formation of invadopodia and invasion by MDA-MB-231 breast adenocarcinoma cells [26]. Indeed, both mDia1 and mDia2 localize to filopodial tips in various mammalian cell lines including tumor cells [27-31]. Previously, mDia1 depletion inhibited Src accumulation at focal adhesions, with effects on adhesion stability as well as cellular migration [32, 33]. Furthermore, mDia2 specifically regulated epithelial cell migration by localizing to the lamella of migrating epithelial cells to polymerize and maintain cortical actin [34]. However, suppression of both intrinsic and direction migration of U87 glioblastoma cells from spheroids in an *ex vivo* model was seen upon mDia agonism [35]. In accordance with these findings, deletion of the *DIAPH3* locus (encoding mDia2) was associated with metastatic disease through its regulation of amoeboid migration in prostate cancer [32, 36]. Indeed, mDia2 was shown to be important for maintaining membrane integrity, as its inhibition by Dia-interacting protein (DIP) led to membrane blebbing and amoeboid motility [38]. In the context of ovarian diseases, disruption of other formins (*e.g.*, mDia3) was associated with effects on ovarian development and premature ovarian failure [32, 39, 40]. Together, these findings indicate a key role for mDia in metastatic processes. However, the mechanism(s) behind invasive transitions in EOC and how it relates to mDia activity and/or localization remains unclear.

In ovarian cancer, decreased E-cadherin expression is associated with invasive peritoneal seeding and is more commonly observed in borderline ovarian tumors and carcinomas compared to benign tumors [43]. Previously, mDia2 depletion promoted ovarian cancer spheroid invasion by driving single cell invasive egress from the spheroid [44]. The exact mechanisms of action remained elusive, however. Here, we show that mDia2 is important for AJ formation and stability in EOC monolayers. Depletion of mDia2 leads to loss of junctional continuity and decreased resistance to mechanical shearing. We observe interactions between mDia2 and α- and β-catenins. Interestingly, mDia2 does not interact with either E- or N-cadherin, indicating that it may not directly interact with the classical junctional cadherin complex. Finally, we show that the interactions between mDia2 and the catenins may not be dependent upon an intact F-actin network. Collectively, these data indicate a critical role for mDia formins in regulating EOC AJs during invasive transitions.

## Results

### mDia2 is essential for junction integrity in spheroids

A key step in EOC dissemination is the shedding of cancer cells from the primary tumor or secondary metastases into the peritoneal cavity, a process that depends on the disruption of cell-cell junctions. To investigate the role of mDia2 in EOC junction integrity, stable mDia2 shRNA OVCA429 cells co-expressing GFP were generated, along with control shRNA GFP-expressing OVCA429 cells. Knockdown (KD) was confirmed with Western blotting (Fig. 1A). E-cadherin expression was unchanged upon mDia2 depletion.

**Figure 1.**
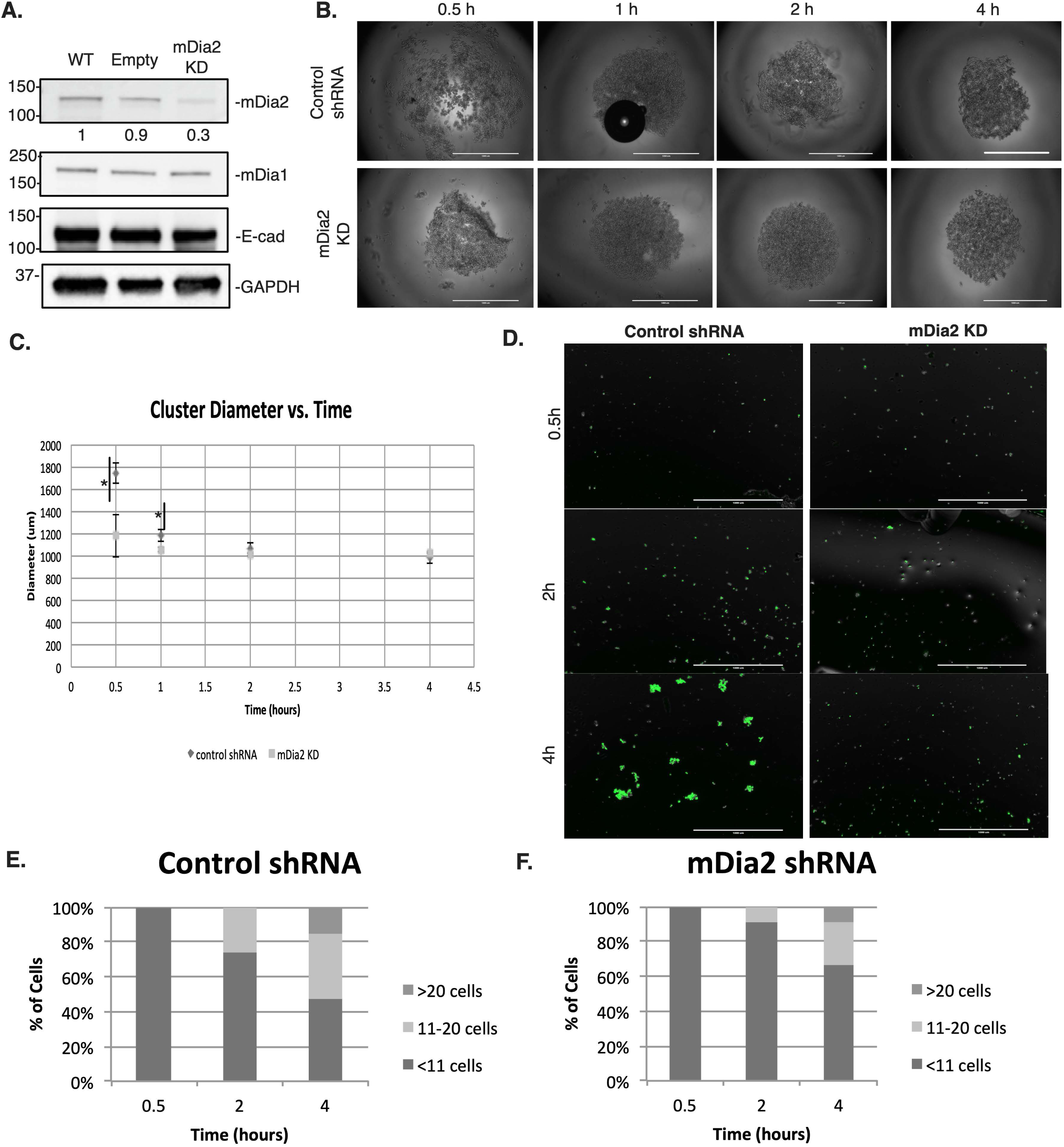
Analysis of mDia2 in a functional cell-cell adhesion assay. **A**. Western blotting for mDia2 in OVCA429-mDia2 KD cells and control knockdown OVCA429 cells. **B**. OVCA429-mDia2 KD and OVCA429-control cells were plated in hanging drop cultures in RPMI media and triturated at specified time points. Representative fields at 0.5, 2, and 4 hours before and after trituration are shown. Scale bar = 1000 μm. **C-D**. Graphs show percentage of cells (OVCA429-mDia2 KD and OVCA429-control) in aggregates of 0-10 cells, 11-20 cells, and >20 cells at specified time points. For each time point, >200 cells were scored. Experiment was performed 3 times in triplicates. A representative experiment of three is shown.

To determine the effect of mDia2 on junctional strength, a hanging drop assay was used to measure cellular resistance to mechanical shear forces [49, 50]. Single-cell suspensions of OVCA429 GFP-expressing mDia2 KD or control cells were seeded as droplets and cultured for 30 minutes to 4 hours (Fig. 1B). Initially at 30 minutes and 1 hour, control cells formed larger and more loosely-packed spheroids compared to mDia2 KD cells. This is consistent with previous findings indicating that mDia2 depletion in ovarian cancer cells led to increased spheroid compaction [44]. However, by 2 hours, mDia2 KD and control cells formed similar sized spheroids (Fig. 1C). At the specified time points, mechanical trituration as applied to the spheroids formed. Cell clusters were then enumerated as either >20 cell, 11-20 cells or <11 cells. OVCA429 mDia2-depleted spheroids were less resistant to shear forces than control cells, as the numbers of clusters >20 cells were reduced to 9% from 16%, respectively, in 4hrs (Fig 1D-F). This corresponded to decreases in cell clusters of 11-20 cells of 26% of OVCA429-control cells compared to 9% of OVCA429-mDia2 KD at 2 hours. These results suggest that mDia2 may have a role in stabilization of cell-cell junctions in EOC clusters and resistance to shear forces.

### Role of mDia2 in AJ formation

In AJ formation, the linkage between cadherins on adjacent cells is dependent on calcium [9]. To determine the effects of mDia2 on AJ formation, a calcium switch assay combined with immunofluorescence (IF) was used to visualize E-cadherin and F-actin in mDia2 KD and control cells. Both mDia2 KD cells and control cells formed AJs when cultured in calcium-containing medium (Fig. 2A). Upon culturing in calcium-free media, AJs in both the mDia2 KD and control cells were abolished, but by 4 hours of calcium repletion with calcium-containing media, AJs were beginning to form in both cells (Fig. 2A). Junctional continuity was then quantified, where a junction was determined to be continuous if the longest E-cadherin-positive region was at least 50% the length between two vertices of a cell-cell junction [16]. The number of continuous E-cadherin-positive junctions was significantly greater for control cells (59% of total junctions) compared to mDia2 KD cells (19% of total junctions) (Fig. 2B). These data indicate that mDia2 may not only have an effect in junction integrity, but also in the formation of the AJ in EOC cells.

**Figure 2.**
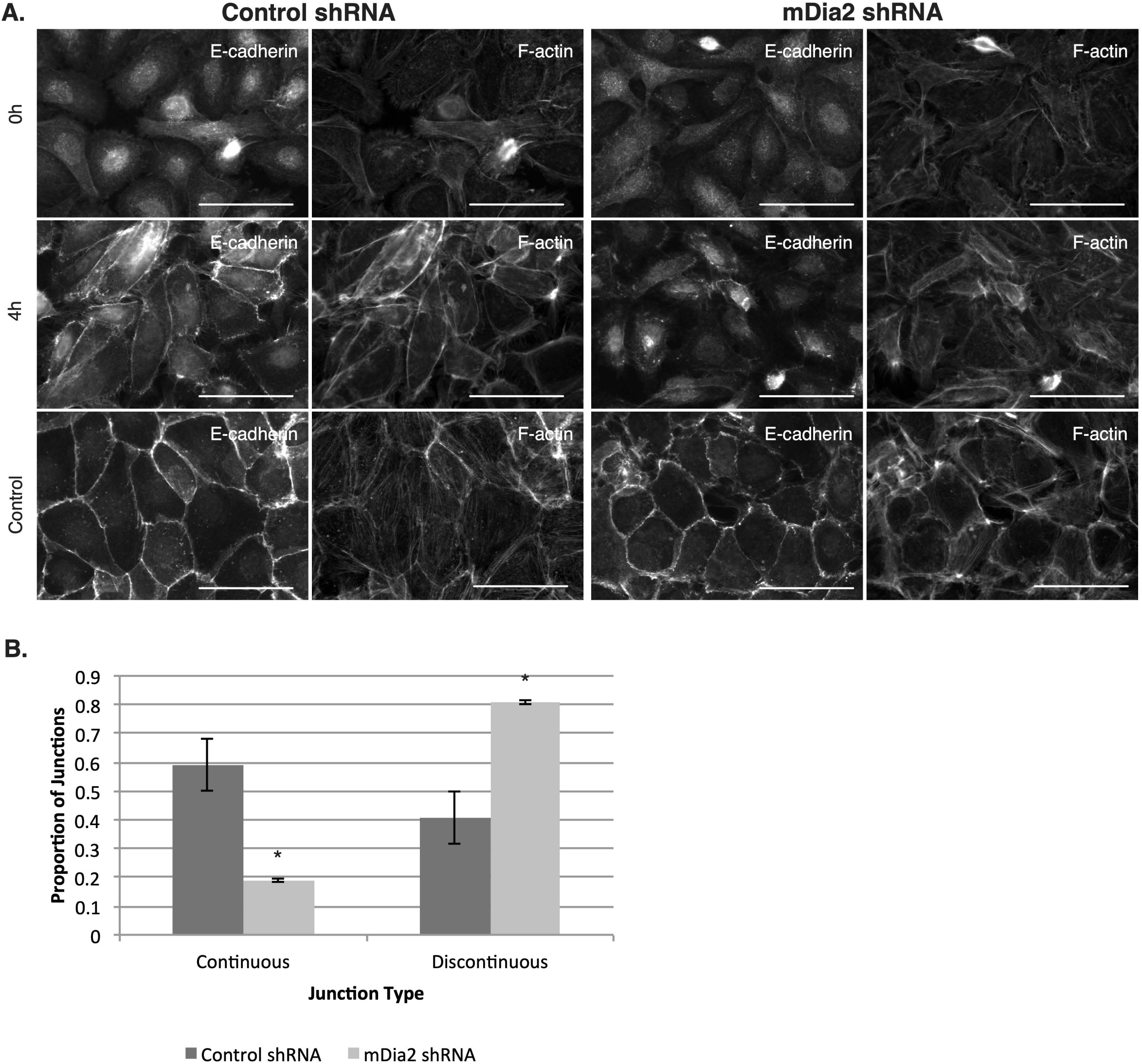
Effect of mDia2 on Adherens Junction formation. **A**. OVCA429-mDia2 KD and OVCA429-control cells were plated and grown to 70% confluence in RPMI medium. Cells were then cultured in calcium-free RPMI for 16 hours followed by regular RPMI and processed and imaged at specified times. For control, cells were in fresh RPMI for 16 hours and again for 4 hours. Cells were stained for E-cadherin and F-actin. Representative fields at 0 and 4 hours after calcium repletion and control cells are shown. Scale bar = 50 μm. **B.** Graphs show increase in proportion of continuous and discontinuous junctions, as measured by E-cadherin stain, in OVCA429-mDia2 KD and OVCA429-control cells from 0 to 4 hours of calcium repletion. A representative experiment of two is shown. *p<0.05. Error bars denote SEM.

### mDia2 interacts with β- and α-catenin but not E-cadherin

As downstream effectors of Rho-GTPases, various formin family members including FMNL2, Formin-1, and mDia1 localized to cadherin-based cell-cell junctions and interact with junctional proteins in epithelial cells [11, 16, 38, 51-55]. Given its effect on AJ formation and stability, we next asked whether endogenous mDia2 interacts with junctional proteins in OVCA429 monolayers. OVCA429 cells express mDia2, mDia1, E-cadherin, N-cadherin, β-catenin, and α-catenin (Fig. 3A-C). Interaction between mDia2 with both α- and β-catenin was detected by co-immunoprecipitation (IP). Previously, Rac1-activated FMNL2 was shown to bind to α-catenin and E-cadherin in MCF10A cells, while Formin-1 was shown to bind to α-catenin in keratinocytes [11, 52, 53]. In our system, mDia2 co-precipitates with both α- and β-catenins, yet does not interact with either E- or N-cadherin, suggesting that mDia2 may be interacting with a cytosolic, rather than membrane-associated junctional pool of catenins (Fig. 3A). mDia1 does not interact with β-catenin (Fig. 3B), indicating a formin-specificity to this interaction. This is consistent with previous findings suggesting that mDia2, and not mDia1, is involved in junctional stability in ovarian cancer spheroids [44]. Neither mDia1 nor mDia2 precipitated with the cadherins (Fig. 3A, C) and Proximity Ligation Assays (PLA) confirmed a lack of mDia2 and E-cadherin interaction in cells (Fig. S1). A positive PLA signal indicates when two proteins are within 40 nm of each other. Collectively, these data indicate that neither mDia1, nor mDia2 associate with AJ-associated E-cadherin complexes.

**Figure 3.**
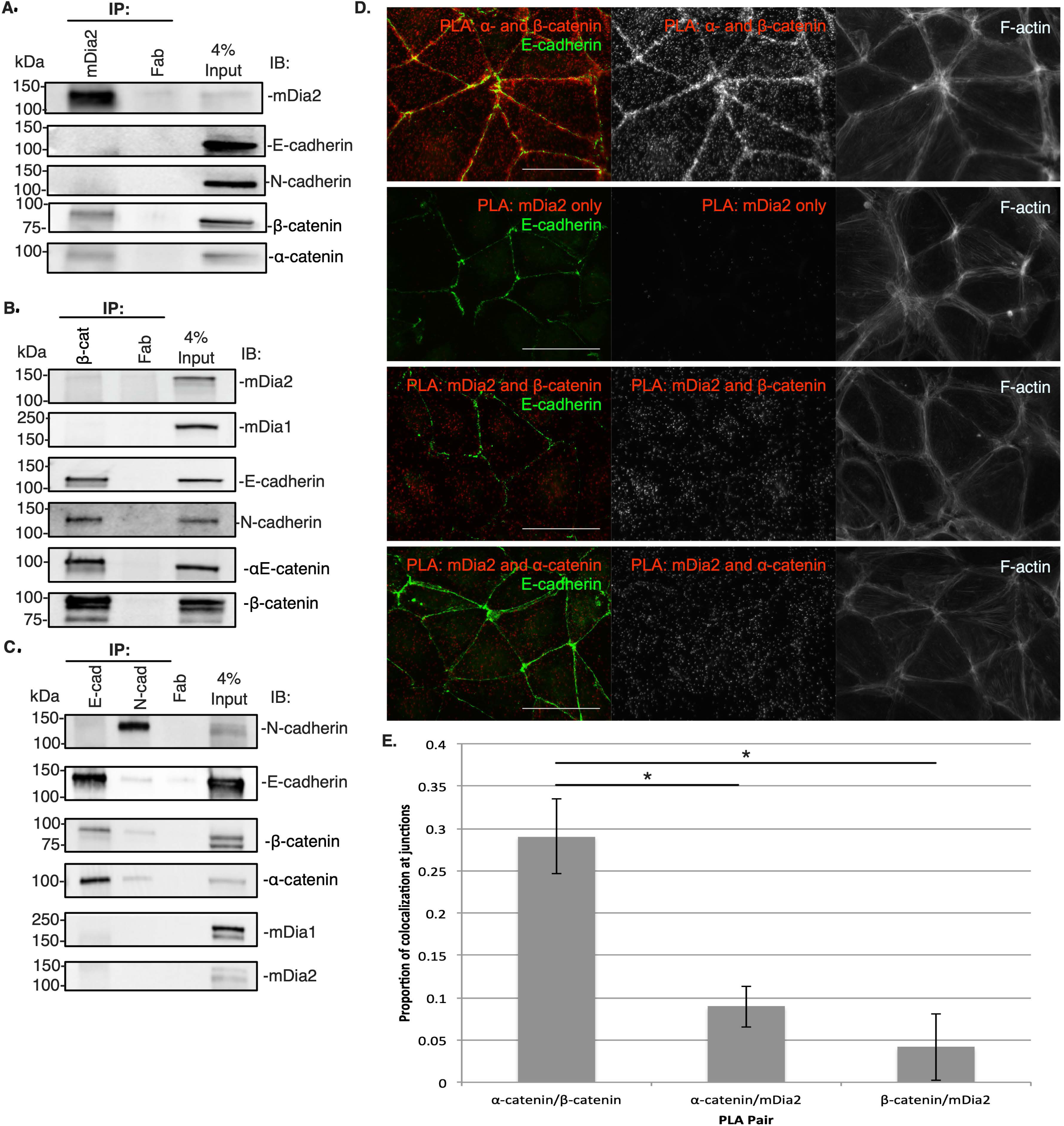
mDia2 interacts with β-catenin and αE-catenin but not E-cadherin in OVCA429 cells. **A.** Immunoprecipitation for mDia2 followed by immunoblotting for mDia2, E-cadherin, β-catenin, and αE-catenin. **B.** IP for β−catenin (β-cat) followed by immunoblotting for mDia2, mDia1, E-cadherin, N-cadherin, β-catenin and αE-catenin. **C**. IP for E-cadherin or N-cadherin followed by immunoblotting for mDia1, mDia2, β-catenin, αE-catenin, E-cadherin and N-cadherin. **A-B** were repeated thrice and representative experiments shown. **D.** Cells were fixed and incubated with primary antibodies against the indicated PLA pairs and E-cadherin. F-actin was stained with phalloidin. Scale bar = 50 μm. **E.** Quantification of junctional mDia2/catenin interactions in **D**. *p<0.01 relative to αE-catenin/β-catenin PLA pair.

To confirm interactions between mDia2 and α- and β-catenins and to localize these interactions within cells, we performed PLA in conjunction with E-cadherin and F-actin IF. PLA interaction between a known direct interaction pair of α- and β-catenin robustly detected both cytosolic and junctional interactions, while PLA reactions with mDia2 antibody alone revealed no signal (Fig. 3D). mDia2 and α- and β-catenin interactions were observed and were mostly localized not to the junctional area as demarcated by E-cadherin, but to the non-junctional area (Fig. 3D, E). Taken together, these results indicate that mDia2 interacts with the α- and β-catenins in a spatially distinct manner than that of junctional E-cadherin.

### mDia2 co-precipitates with α- and β-catenin in HEK293 cells

To validate the interactions between mDia2 and α- and β-catenins, we exogenously expressed mDia2 and α- or β-catenin in HEK293 cells. IP for GFP revealed GFP-mDia2 interaction with HA-α-catenin (Fig. 4A). Next, we transfected HEK293 cells with Flag-mDia2 and GFP-β-catenin. IP for GFP showed GFP-β-catenin interacted with Flag-mDia2 (Fig. 4B). These results confirm our previous findings in OVCA429 cells, confirming interactions between mDia2 and α- and β-catenin.

**Figure 4.**
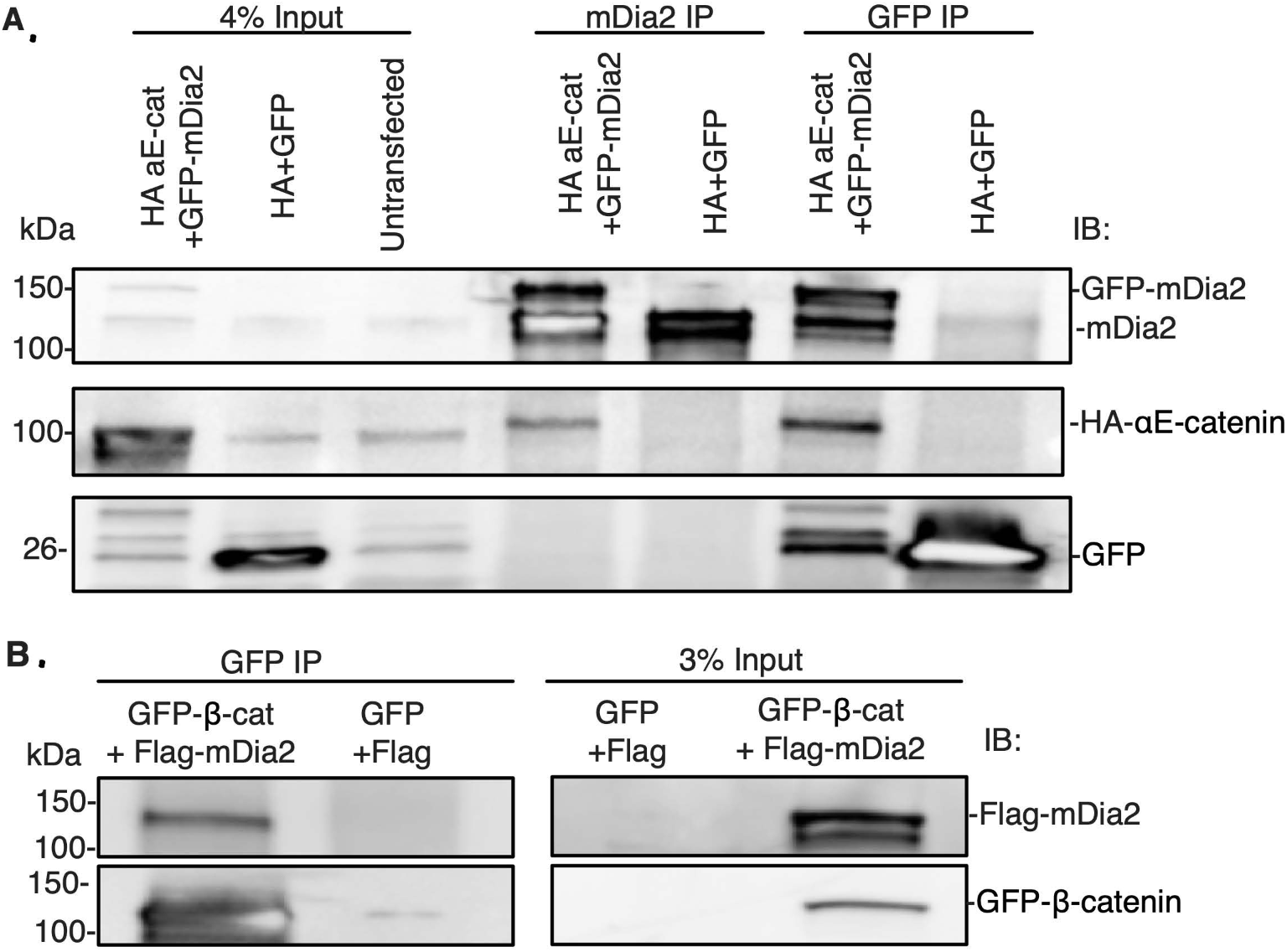
mDia2 co-precipitates with αE-catenin in HEK293 cells. **A**. Cells were transfected with GFP-mDia2 and HA-α-catenin or GFP and HA empty vectors. IP for GFP was followed by immunoblotting for mDia2, α-catenin, and GFP. The experiment was repeated thrice. **B**. Cells are transfected with Flag-mDia2 and GFP-β-catenin or Flag and GFP empty vectors. IP for GFP was followed by immunoblotting for Flag and GFP.

### mDia2 affects junctional stability

To assess the requirement for mDia2 on junctional protein expression and localization, we visualized junctional markers E-cadherin and β-catenin in mDia2- and control-depleted OVCA429 cells. While total E-cadherin levels were not affected by mDia2 depletion relative to control cells (Fig. 1A), mDia2 knockdown cells exhibited clustered and discontinuous localization of E-cadherin and β-catenin at the junctional region compared to the linear junctional staining of both proteins in control cells (Fig. 5A). This effect was similar to the clustered staining of E-cadherin and β-catenin seen upon treatment with untransfected OVCA429 cells with a small molecule inhibitor of the formin homology 2 (FH2) domain (SMIFH2), a broad-spectrum formin inhibitor (Fig. S2A) [56]. The number of continuous junctions was quantified. The proportion of continuous junctions was significantly higher for control cells compared to mDia2 knockdown cells. In control cells, 76% of junctions between cells were continuous, compared to 22% of mDia2 knockdown cell junctions (Fig. 5B). This indicates that mDia2 is essential for junctional localization of E-cadherin and β-catenin, and is consistent with our previous findings suggesting loss of junction strength upon mDia2 depletion.

**Figure 5.**
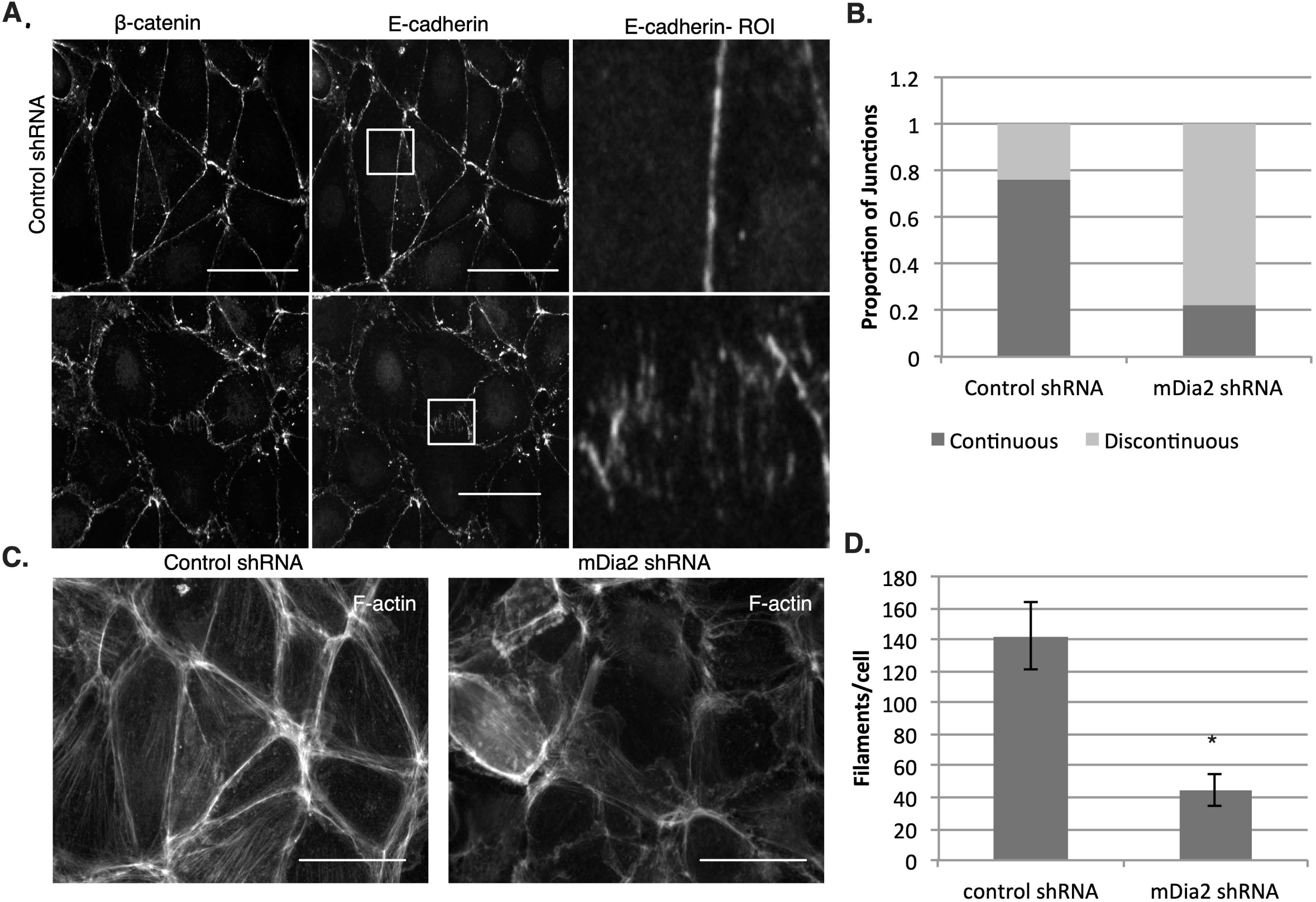
mDia2 expression affects junctional stability. **A**. OVCA429-mDia2 KD and OVCA429-control cell monolayers were stained for E-cadherin and β-catenin. Coverslips were imaged with fluorescence microscopy at 60x magnification. Boxes denote regions of interest (ROI). Scale bars = 50 μm. **B**. Graphs show ratios of continuous to discontinuous adherens junctions as measured by E-cadherin staining. Three fields and at least 50 junctions per condition were quantified. **C.** Phalloidin was used to stain for F-actin in OVCA429-mDia2 KD and OVCA429-control cells. Scale bars = 50 μm. **D**. Actin filaments per cell for OVCA429-mDia2 KD and OVCA429-control cells are shown. Five separate fields and at least 70 cells per condition were quantified. The experiment was repeated thrice. *p<0.05.

To evaluate F-actin levels underlying AJs in mDia2- and control-depleted cells, we stained OVCA429s with phalloidin. We observed a marked decrease in F-actin staining in the mDia2-depleted cells relative to control cells (Fig. 5C). Actin filaments were quantified using Filaquant software, which showed a significant reduction in F-actin in mDia2 knockdown cells relative to control cells (Fig. 5D), just as formin inhibition with SMIFH2 also reduced filament levels (Fig. S2B). These results suggest that mDia2 depletion leads to disruption in junctional localization of β-catenin and E-cadherin with concurrent reduction in F-actin.

### mDia2 expression affects interactions between junctional proteins

To determine whether a causal relationship exists between F-actin reduction and AJ disruption, we investigated interactions between junctional proteins upon mDia2 depletion. AJs are characterized by stable interactions between E-cadherin and catenins (reviewed in [57]). We therefore evaluated the interactions between E-cadherin and β-catenin in mDia2 knockdown and control OVCA429s. We used PLA probes to visualize E-cadherin and β-catenin interaction (Figure 6A), as well as α-catenin and β-catenin (Figure 6B), in conjunction with phalloidin staining. We observed a decrease in β-catenin/E-cadherin interaction in mDia2 knockdown cells with notably more disorganized F-actin (Fig. 6A). Quantification revealed a significant reduction in β-catenin/E-cadherin interaction in mDia2 knockdown cells compared to control cells (Fig. 6C). This indicates that mDia2-mediated junction stability may, in part, be attributed to an F-actin-dependent stabilization of the β-catenin/E-cadherin complex.

**Figure 6.**
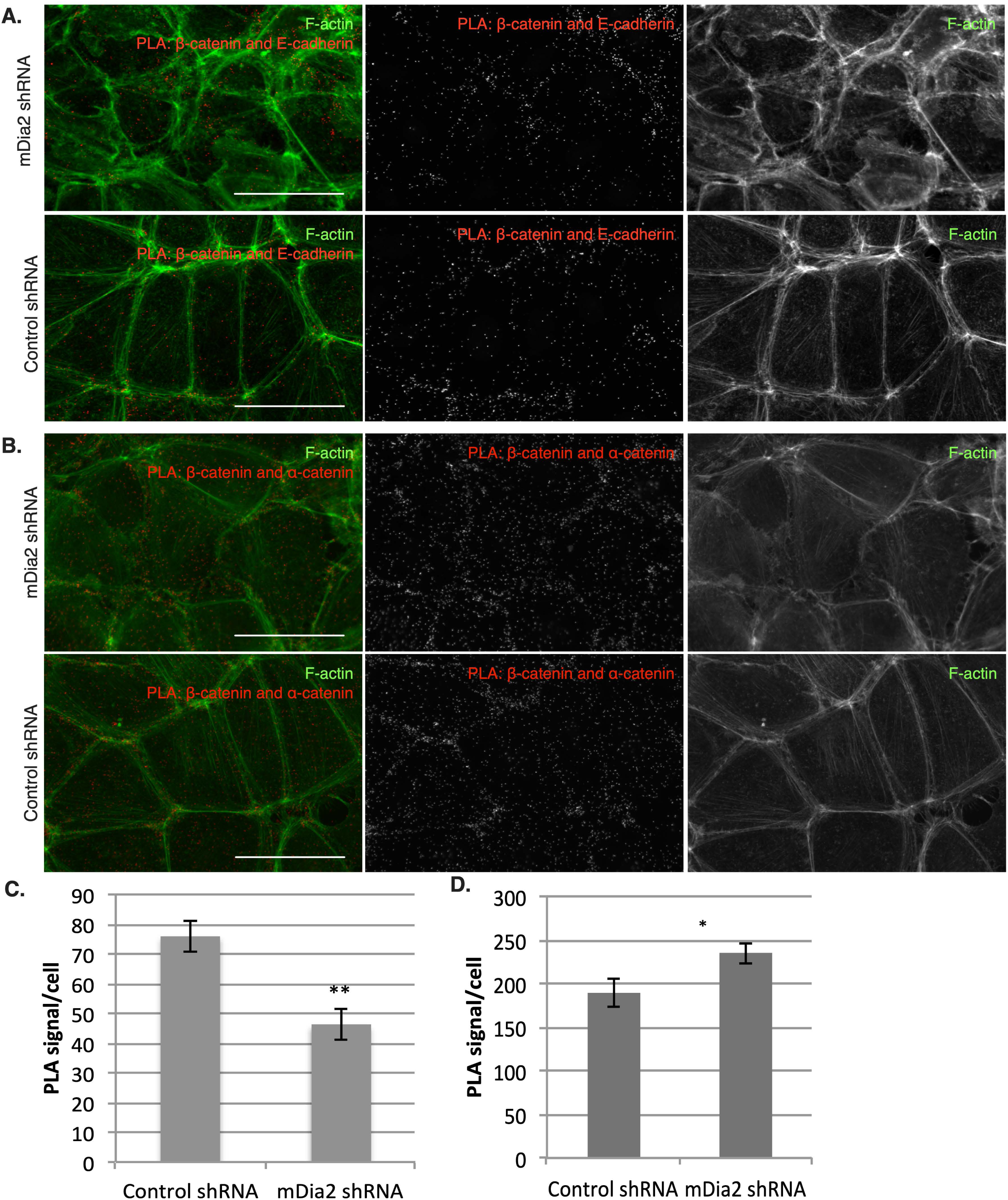
mDia2 expression affects interactions between junctional proteins. **A**. Representative images of OVCA429-mDia2 KD and OVCA429-control cells labeled to visualize F-actin and β-catenin/E-cadherin interactions by PLA. **B.** Quantification of PLA detecting β-catenin/E-cadherin interactions (n=60, 101). **C.** Representative images of OVCA429-mDia2 KD and OVCA429-control cells labeled to visualize F-actin and α-catenin/β-catenin interactions by PLA. **D.** Quantification of PLA detecting α−catenin/β-catenin interactions. Six fields were analyzed per condition **(B**,**D).** *p<0.05, **p<0.001. Scale bars = 50 μm. All values denote mean +/- SEM.

As β-catenin is also associated with both junctional and cytosolic α-catenin, we investigated whether mDia2 depletion impacts α- and β-catenin interactions using PLA probes to visualize α- and β-catenin interactions in conjunction with F-actin. Interestingly there was a significant increase in α- or β-catenin interactions in mDia2 knockdown cells compared to control cells (Fig. 6B, D). These results suggest that mDia2 promotes E-cadherin/β-catenin interactions while preventing α-/β-catenin interactions. Whether one interaction occurs at the expense of the other is unclear as both α- and β-catenin and their interacting complex have multiple effects on the cell, including AJ stabilization and transcriptional regulation.

### Actin disruption does not inhibit interactions between mDia2 and α- or β-catenin

We previously showed a reduction in levels and organization of F-actin in mDia2 knockdown OVCA429 cells (Fig. 5C-D). This was in response to global suppression of mDia2-dependent F-actin dynamics. We therefore evaluated whether F-actin was necessary to facilitate interactions between mDia2 and α- or β-catenin or between β- and α-catenin. We used cytochalasin D (CytoD) to globally inhibit actin polymerization. PLA was used to visualize interactions between these protein pairs in CytoD- or DMSO-treated OVCA429s. By 30 minutes, CytoD-treated cells had visible and widespread disruption in the cytoskeletal network accompanied by AJ disruption as visualized by E-cadherin staining (Fig. 7A-C). Cortical actin underlying the junctions was clustered and discontinuous in CytoD-treated cells compared to the continuous linear junctional staining seen in control cells. Interestingly, there was comparable PLA signal for the α-/β-catenin and β-catenin/mDia2 PLA pairs between treatment and control cells (Fig. 7A, C-D), suggesting that the interaction between mDia2 and β-catenin or α− and β-catenin do not occur through mutual binding to F-actin. This is consistent with previous studies, which pointed to an α-catenin pool that heterodimerizes with β-catenin and concurrently displays decreased affinity for F-actin [13, 14]. This pool of α-catenin should be unaffected by disturbances in the actin cytoskeleton. Interestingly, a small yet significant increase in mDia2/α-catenin interactions was observed upon CytoD treatment and global F-actin polymerization defects (Fig. 7B, D). These data collectively suggest that mDia2 can interact with α-catenin in the absence of organized F-actin networks. Indeed, previously Formin-1 was shown to interact with α-catenin in a purified system in absence of F-actin [53]. However the primary α-catenin-binding sequence of Formin-1 is not significantly homologous to that of mDia formins, suggesting an alternate mechanism. Furthermore, suppression of polymerized F-actin may underlie conformational changes in α-catenin to enhance its localized interaction with mDia2 away from the AJ [8, 12, 58]. Together, this suggests that actin disruption does not prevent interactions between mDia2 and the catenins, but instead may alter the nature and location of these interactions, such that they are no longer contribute to junctional stabilization.

**Figure 7.**
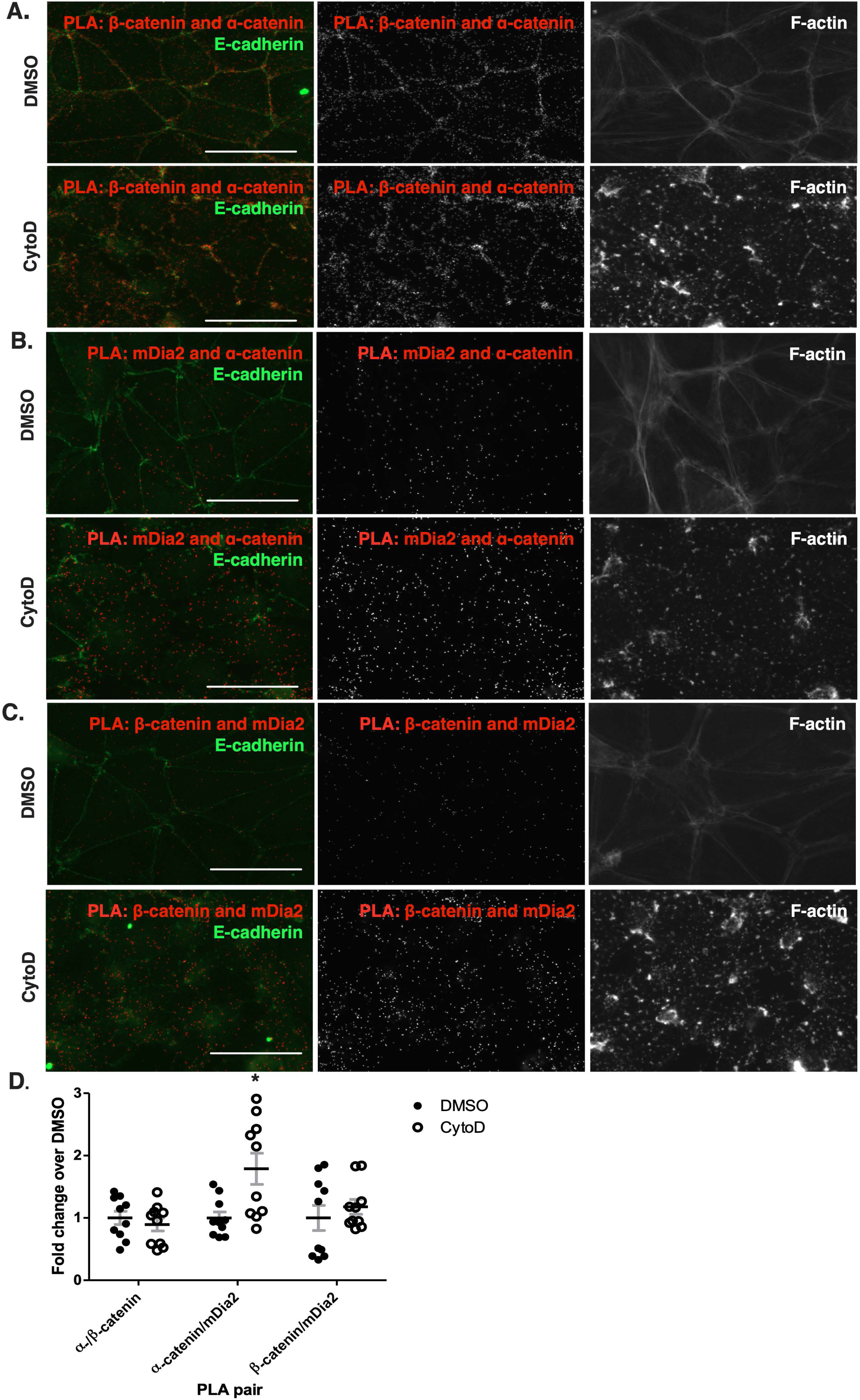
Actin disruption does not inhibit Interactions between mDia2 and α- or β-catenin. **A-C**. Representative images of OVCA429-mDia2 KD and OVCA429-control cells labeled to visualize α-catenin/β-catenin, α-catenin/mDia2, or mDia2/β-catenin by PLA are shown. **D**. Quantifications of PLA detecting α-catenin/β-catenin, α-catenin/mDia2, or mDia2/β-catenin interactions. Ten fields analyzed per condition. All values denote mean+/-SEM. *p<0.05.

## Discussion

The poor prognosis associated with EOC can largely be attributed to its diagnosis in the late stages of the disease when cancer cells have already disseminated within the peritoneal cavity. Previously, depletion of mDia2, but not mDia1, was associated with single cell invasion from ovarian cancer spheroids [44]. Here, we show for the first time that mDia2 is essential for maintenance of cell-cell junction strength in EOC spheroids and AJ formation. This may be attributed to interactions between mDia2 and AJ proteins, specifically α- and β-catenin. Interestingly, mDia2 does not appear to interact with the catenins at the junctions, indicating that its localization to the AJ is not necessary for its role in junction maintenance. As expected, mDia2 depletion resulted in a significant reduction in actin filament levels and marked disorganization in cytoskeleton architecture. However, disruption of the F-actin network did not prevent interactions between α- and β-catenin or between mDia2 and β-catenin. These data suggest a key role for mDia2 in AJ formation and stability in EOC cells which may not be entirely dependent on its actin-polymerizing activity. In this study, we did not assess the roles of targeting mDia2-directed microtubule stabilization in AJ function, as mDia formins were shown to strengthen AJs in the absence of microtubules [16].

Epithelial cells, including EOC cells, develop cadherin-based AJs, via a three-step process which involves initiation of the cell-cell contact, expansion of the contact interface, and finally, stabilization of the contact to form multi-cellular structures including epithelial monolayers and three-dimensional (3D) spheroids [11, 57, 59, 60]. During cell migration, actin polymerization generates dynamic plasma-membrane protrusions, which include finger-like filopodia characterized by parallel F-actin bundles extending beyond the leading edge of lamellipodia [60, 61]. Contact initiation occurs when junctional proteins at the filopodia tips form homophilic linkages with proteins in the neighboring cell [9, 11, 62]. Cadherins then cluster at the newly formed contact and induce actin-remodeling to form a stable junction [11, 59, 60]. Our findings that mDia2 plays a role in junction formation are consistent with role for other formins including Formin-1 and mDia1 in junction stabilization in keratinocytes and MCF7 breast cancer cells [11, 16, 53]. This, compounded with the finding that mDia2 is a predominant formin regulating ovarian cancer spheroid organization [44] supports a key role for mDia2 in AJ formation and stabilization in EOC.

Stability of AJs is thought to depend upon anchorage of the junctional complex of E-cadherin and catenins to the underlying F-actin cytoskeleton [10]. Traditional models have implicated α-catenin as the link between the E-cadherin/β-catenin junctional complex and the underlying F-actin [13, 14]. Yet, binding to β-catenin decreased the affinity of α-catenin for F-actin, so an alternative model was proposed wherein α-catenin cycles between a junctional pool bound to the E-cadherin/β-catenin complex and a peri-junctional pool bound to F-actin along with various actin-binding proteins such as the formins [8, 12, 13]. In the present study, mDia2 associated with both α- and β-catenins, but neither with E-nor N-cadherin. Furthermore, these mDia2-catenin interactions occurred predominantly in the non-junctional region. This suggests that mDia2 either associates with α- and β-catenin separately as a duplex or together as a cytosolic or nuclear triplex. We cannot rule out the possibility of a triple complex of E-cadherin, β-catenin, and mDia2, although our co-IPs failed to support this notion. At the same time, there remains the possibility that under certain specific conditions, mDia2 may act as the link between the E-cadherin complex and α-catenin bound to F-actin.

Formins are known to maintain junctional stability through both cortical actin polymerization and junctional contractility regulation. Indeed, mDia1 regulates junctional tension by reorganizing actin into actomyosin bundles at the AJ in Caco-2 colon epithelial cells [64]. This is significant given that α-catenin binding to both F-actin and the actin-binding protein vinculin are force-dependent due to conformational changes in α-catenin that occur with force application [6, 65]. As mDia2 is important for junctional stability and resistance to shear force in EOC cells (Figure 1), it is reasonable to surmise that mDia2 stabilizes junctions by providing contractile force at the AJ. This, in turn, would enhance anchorage of the junctional E-cadherin/β-catenin complex to the F-actin network via α-catenin.

What if the actin network itself is disrupted? While inhibition of actin polymerization with CytoD led to AJ disruption, it did not reduce non-junctional mDia2 and β-catenin interactions (Fig. 7C-D). Meanwhile, F-actin disruption slightly increased interactions between mDia2 and α-catenin (Fig. 7B, D). Therefore, although AJ stability involves actin-dependent localization of junctional proteins [66], mDia2 does not require F-actin to associate with either α- or β-catenin in the non-junctional region. As β-catenin is not known to bind to F-actin, actin depolymerization is not expected to affect its interaction with mDia2. While α-catenin homodimers bind to F-actin in the cytosol, α-catenin’s major stoichiometric binding partner in the cytosol is β-catenin [12, 67]. The slight increase in α-catenin interaction with mDia2 could potentially be attributed to a change in α-catenin conformation that occurs upon actin depolymerization, just as α-catenin conformation is regulated by actomyosin contractile force [64].

We show here that mDia2 depletion significantly reduces global F-actin levels (Fig. 5D). Decreased F-actin could potentially decrease the number of α-catenin homodimers that preferentially binds to it, concomitantly increasing the pool of α-/β-catenin heterodimers. Indeed, we observed that mDia2 depletion is associated with an increase in α-/β-catenin interaction. We also observed a concurrent reduction in β-catenin/E-cadherin interaction. This is consistent with the concept that stability of the junctional β-catenin/E-cadherin complex is dependent on its anchorage to F-actin. Together, these findings support the notion of mDia2’s indirect role in junction stabilization through F-actin polymerization and bundling.

Our findings are consistent with recent publications that propose a role for formins in regulating the epithelial mesenchymal transition (EMT). Both formin inhibition with SMIFH2 and depletion of mDia1 and mDia2 prevented TGF-β-induced EMT in lung, mammary, and renal epithelial cells [68]. Others demonstrated the role of formins including FHOD1 and FMNL2 in the morphological changes associated with EMT [68-70]. In ovarian cancer, decreased E-cadherin expression is associated with peritoneal seeding of tumor cells and lower overall survival rate [43, 71]. Here we identify a novel role for mDia2 in AJ stabilization to impact EMT in 3D ovarian cancer spheroids [44]. While we did not observe an interaction between mDia2 and the cadherin molecules, mDia2 interacts with key regulators of the AJ, α- and β-catenin to disrupt E-cadherin localization to AJs.

It is interesting to note that mDia2 minimally interacts with junctional β-catenin, which was unexpected given its junction stabilizing effect. It has previously been proposed that the AJ acts as a sink for cytosolic β-catenin, drawing β-catenin away from the Wnt signaling pathway with transcriptional activation of pro-migratory genes [72]. Cytosolic and nuclear α-catenin can also regulate transcription, both through β-catenin binding and its regulation of nuclear actin [73]. Our finding that mDia2 interacts with both catenins may suggest a potential role in transcriptional regulation, bridging the AJ and Wnt signaling pathways. Future studies would aim to determine how mDia2 affects Wnt/β-catenin signaling in EOC.

## Conclusions

In summary, our findings indicate an essential role for mDia2 in AJ formation and stability in EOC cells. These effects are likely achieved through its interactions with and regulation of α- and β-catenin. While we demonstrate interaction, it remains uncertain whether α-catenin binding to mDia2 impacts either actin polymerization or bundling, or which domains of mDia2 and α-catenin interact. Furthermore, our current studies utilize EOC monolayers to dissect the interactions between mDia2 and proteins involved in the AJ. Assuming that these same interactions occur in 3D spheroids and given that loss of mDia2 is associated with disease progression in ovarian cancer [74], our findings support a novel mechanism for EOC dissemination that should be considered in development of targeted therapy against this deadly disease.

## Methods

### Cell lines and reagents

Serous ovarian adenocarcinoma OVCA429 cells were kind gifts from Dr. Deborah Vestal (University of Toledo, Toledo, OH) were grown in RPMI-1640 (GE Lifesciences (Pittsburgh, PA)) containing 10% (v/v) fetal bovine serum (FBS), 100 U/ml penicillin, and 100 μg streptomycin. HEK293 human embryonic kidney cells were from ATCC (Manassas, VA) and were grown in DMEM (GE Lifesciences) containing 10% (v/v) FBS, 100 U/ml, and 100 μg streptomycin. All cells were grown in a 37°C incubator with 5% CO_2_.

OVCA429 cells were plated at 200,000 cells per 35 mm well, grown to 70-80% confluence, then treated with DMSO or 40 μM SMIFH2 in DMSO (EMD Biochemicals, Tocris Bioscience, Avonmouth) in full media for 8 hours.

### Western blotting

Cells were harvested for Western blots using SDS lysis buffer (0.5 M Tris-HCl, pH 6.8, glycerol, 10% SDS (wt/vol), 0.1% bromophenol blue (wt/vol), 0.1 M diothiothreitol (DTT)). Lysates were separated using 4-20% gradient SDS-PAGE gels (BioRad, Hercules, CA) and were transferred to PVDF membranes using the BioRad Trans-Blot turbo system.

### Transfection and knockdown

pCMV-driven plasmid vectors encoding GFP, GFP-mDia2 as well as Flag-mDia2 were kind gifts of Dr. Art Alberts (Van Andel Institute, Grand Rapids, MI). Plasmids encoding HA and HA-α−catenin were kind gifts of Dr. Deniz Toksoz (Tufts University, Medford, MA). GFP-β-catenin was from Addgene (Cambridge, MA) and GFP empty vector was a kind gift from Dr. Kam Yeung (University of Toledo).

For knockdown experiments, mDia1 siRNA (J-010347-070005) and control GAPDH (D-001140-01-05) siRNA constructs were purchased from Thermo Scientific (Waltham, MA), and mDia2 shRNA and control pGFPVRS from Origene (Rockville, MD) respectively.

To generate OVCA429 cells stably depleted of mDia2, OVCA429 cells were transfected with GFP-mDia2 shRNA constructs (Origene) using Fugene (Promega (Madison, WI)) per manufacturer’s protocol and drug selected with 4 μg/ml puromycin. Control cells were generated using empty pGFP-V-RS vector (Origene). Cells were then further selected for GFP through flow cytometry using the FACS Aria Ilu High-Speed Cell Sorter (BD Biosciences (Franklin Lakes, NJ)). Knockdown of mDia2 was confirmed using Western blotting with an anti-mDia2 antibody (Proteintech Rosemont, IL) at 1:1000.

HEK293 cells were transiently transfected using a standard calcium phosphate transfection method [45].

### Immunoprecipitation

OVCA429 cells were grown to 70-80% confluence and serum starved overnight with RPMI-1640 containing 0.1% (v/v) FBS and 100 μg streptomycin. Cells were then serum stimulated with RPMI-1640 containing 10% (v/v) FBS and 100 μg streptomycin for 4 hours. Lysates were collected with NP40 lysis buffer (20 mM Tris-HCl pH 7.5, 100 mM NaCl, 1% NP40, 10% glycerol) with protease inhibitors (1 μM each of NaVO_4_, aprotinin, pepstatin, leupeptin, DTT, PMSF). Lysates were incubated with anti-mDia2 antibody (Proteintech (Rosemont, IL)) or control Fab fragment (Jackson Immunolabs) at a concentration of 1 μg antibody per 1 mg lysate for 3 hours at 4°C with shaking, followed by addition of Protein A Agarose beads (Invitrogen, Santa Cruz) for 1 hour at 4°C with shaking. Beads were washed 5 times with NP40 buffer then heated at 85°C in SDS lysis buffer prior to loading in gels.

HEK293 cells were grown to 70% confluence and transfected with HA and GFP or HA-α-catenin and GFP-mDia2-encoding vectors. Lysates were collected with NP40 buffer and 1 μM protease inhibitors 48 hours post-transfection. Lysates were incubated with anti-GFP (Abcam (Cambridge, United Kingdom)) antibody for 3 hours at 4°C then 1 hour with Protein A agarose beads, prior to washing in NP-40 buffer.

Western blotting was performed with 4-20% SDS-PAGE (Biorad), followed by immunoblotting with the following antibodies: rabbit anti-mDia2 (1:1000, (Proteintech), mouse anti-E-cadherin (1:1000, Cell Signaling), mouse anti-N-cadherin (BD Transduction Laboratories (Franklin Lakes, NJ)), rabbit anti-mDia1 (1:1000, (Proteintech), rabbit anti-α-catenin (1:2000 (Proteintech), and rabbit anti-β-catenin (1:2000 (Proteintech) and visualization by the Clarity™ Western ECL kit (Biorad).

### Immunofluorescence and Image Analysis

For immunofluorescence, cells grown upon glass coverslips were fixed in 4% paraformaldehyde (PFA)/phosphate buffered saline (PBS) for 5 minutes, washed with PBS, permeated with 0.5% Triton X-100 (TX100) for 20 minutes, blocked for an hour with 3% bovine serum albumin (BSA)/PBS, and incubated with antibodies against 1:100 β-catenin (Proteintech) and 1:100 E-cadherin (Cell Signaling), or 1:100 αE-catenin (Genetex (Irvine, CA)) and 1:100 mDia2 (Proteintech) overnight at 4°C followed by incubation with 1:200 Alexa-Fluor secondary antibodies (Invitrogen) for 2 hours at 37 °C. To visualize F-actin and nucleus, we used 1:100 Alexa Fluor 647 Phalloidin (ThermoFisher Scientific (Waltham, MA)) and DAPI (Invitrogen), respectively. Coverslips were mounted with Fluoromount-G (SouthernBiotech (Birmingham, AL)) and visualized with an Olympus 60x UPlanFl 1.25 NA oil objective on the EVOS FL epifluorescence microscope (AMG/Thermo Fisher).

To quantify actin filament number and length, images of phalloidin-stained cells were uniformly processed with Photoshop and Filaquant software provided by Dr. Konrad Engel of the University of Rostock [46]. To determine junction continuity, a junction was characterized as “continuous” as described [16]. Briefly, if E-cadherin fluorescence along a cell-cell contact was above background fluorescence for at least 50% of the distance between the cell vertices, that junction was considered continuous [16]. Quantification was performed using a custom Python script. At least 50 junctions were counted per condition for each experiment and the experiment was performed thrice.

### *In Situ* Proximity Ligation Assay (PLA)

To visualize the interactions and localization of the interactions between mDia2 and αE-catenin, we used the Duolink PLA kit (Sigma-Aldrich (St. Louis, MO)). OVCA429 cells were fixed with 4% PFA/PBS, permeated with 0.5% TX100, blocked with 3% BSA/PBS and incubated overnight with primary antibodies. The following antibodies were used: 1:100 goat anti-E-cadherin (R&D Systems (Minneapolis, MN)), 1:100 rabbit anti-mDia2 (Proteintech), 1:100 mouse anti-αE-catenin (GeneTex), 1:100 mouse anti-β-catenin (Origene), 1:100 mouse anti-E-cadherin (Cell Signaling). Cells were then incubated with secondary antibodies conjugated to oligonucleotides per manufacturer’s protocol. Briefly, after incubation with anti-mouse and anti-rabbit secondary antibodies, cells were incubated with ligase followed by polymerase and close protein-protein interactions (<40 nm apart) were detected as fluorescent dots generated by rolling circle amplification with complementary fluorescent oligonucleotides [47]. Cell nuclei were stained with DAPI (Invitrogen) and F-actin with 1:100 Alexa Fluor 647 Phalloidin (ThermoFisher Scientific). Quantification of colocalization was performed using ImageJ Particle Analysis and Colocalization plugin. Three independent fields and at least 47 cells per condition were quantified.

To visualize interactions between mDia2 and α-catenin/β-catenin, or between αE-catenin and β-catenin in OVCA429 cells upon cytochalasin D treatment, cells were plated at 200,000 cells/well into a 6-well dish upon glass coverslips. At 70-80% confluence, cells were treated with 1 μM cytochalasin D (Calbiochem (Burlington, MA)) for 30 minutes, then fixed and stained for PLA pairs (mDia2/α-catenin, mDia2/β-catenin, or α-catenin/β-catenin), E-cadherin, F-actin, and DAPI as above.

### Hanging Drop Assay

The assay was performed as described [48, 49]. Briefly, control or mDia2 KD OVCA429 cells were trypsinized, centrifuged, and re-suspended as single- or up to 3-cell suspensions at 2.5×10^5^ cells/ml. For each cell type, 20-μl droplets were pipetted onto the lids of 35 mm culture dishes and dishes were filled with 2 ml of growth media. At 0.5, 2, and 4 hours, the lids were inverted and drops were transferred to glass slides and pipetted 10 times through a 20-μl pipet tip. Three random fields were imaged using epifluorescence microscopy with an Olympus 4x UPlanFL 0.13 NA objective lens and numbers and sizes of clusters quantified. At least 200 cells were counted per condition. The experiment was performed thrice.

### Calcium Switch Assay

Untransfected, control knockdown, and mDia2 KD OVCA429 cells were grown to 60-70% confluence upon glass coverslips and incubated with RPMI-1640 with 0.1% FBS, 100 U/ml penicillin, and 100 μg streptomycin, without calcium (US Biological (Salem, MA) for 16 hours. Medium was then changed to either the same calcium-free RPMI-1640 or RPMI-1640 with 0.42 mM calcium (GE Lifesciences) and 10% FBS and 100 mg streptomycin for 4 hours. Cells were fixed with 4% PFA/PBS at given time points and stained for E-cadherin and β-catenin. Junction continuity was determined as above. At least 50 junctions were scored per condition at each time point and the experiment was performed twice.

### Statistics

Two-tail Student’s t-tests were used with a 95% confidence value. P-values less than 0.05 were interpreted as statistically significant. All error bars denote standard deviations from representative experiments unless otherwise indicated. Graphs and statistics were generated from Microsoft Excel and GraphPad Prism software.

## Abbreviations

3D: three dimensional
AJ: adherens junction
CytoD: cytochalasin D
EMT: epithelial mesenchymal transition
EOC: epithelial ovarian cancer
FH2: formin homology 2
HEK: human embryonic kidney
IF: immunofluorescence
IP: immunoprecipitation
KD: knockdown
mDia: mammalian Diaphanous
PLA: proximity ligation assay
SMIFH2: small molecule inhibitor of FH2

## Declarations

### Ethics approval and consent to participate

Not applicable

### Consent for publication

Not applicable

### Availability of data and materials

All data generated and analyzed in this study are available upon reasonable request from the corresponding author.

### Authors contributions

Y.Z. and K.M.E. conceived of the experiments. Y.Z. and K.P. performed experiments and analysis, and wrote the manuscript draft. K.M.E. and K.N.B. edited the manuscript draft. All authors approved the final submission.

## Acknowledgements

We thank members of the Eisenmann lab, Drs. Rafael Garcia-Mata, William Maltese, Randall Ruch, Eda Yildirim-Ayan, Andrea Kalinoski, Dayanidhi Raman and Peterson G.T. Schwifty for discussion and technical guidance. We also thank the University of Toledo Department of Cancer Biology for everyone’s support and generosity with their time and resources, especially Nicole Bearss and Augustus Tilley for guidance with PLA experiments. We thank Drs. Sahezeel Awadia and Ashtynn Zinn for technical assistance, Dr. Konrad Engel for use of Filaquant, and Dr. Kam Yeung and Dr. Deniz Toksoz for their valuable time and plasmids.

## Supplemental Figure Legends

**Supplemental Figure 1.**
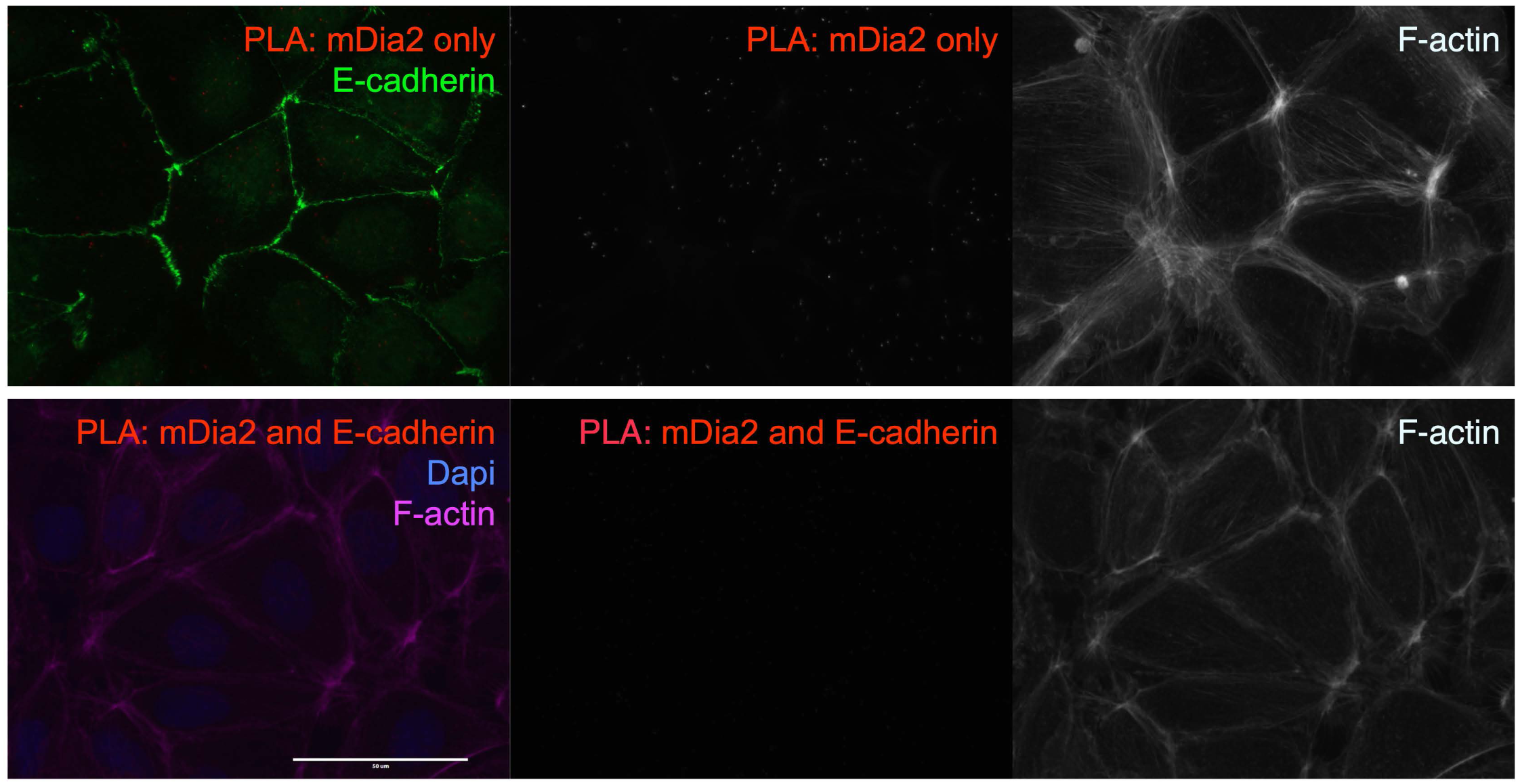
mDia2 does not interact with E-cadherin. OVCA429 cells were treated with PLA probes targeting mDia2 only or mDia2 and E-cadherin. F-actin is stained with phalloidin and nuclei with DAPI.

**Supplemental Figure 2.**
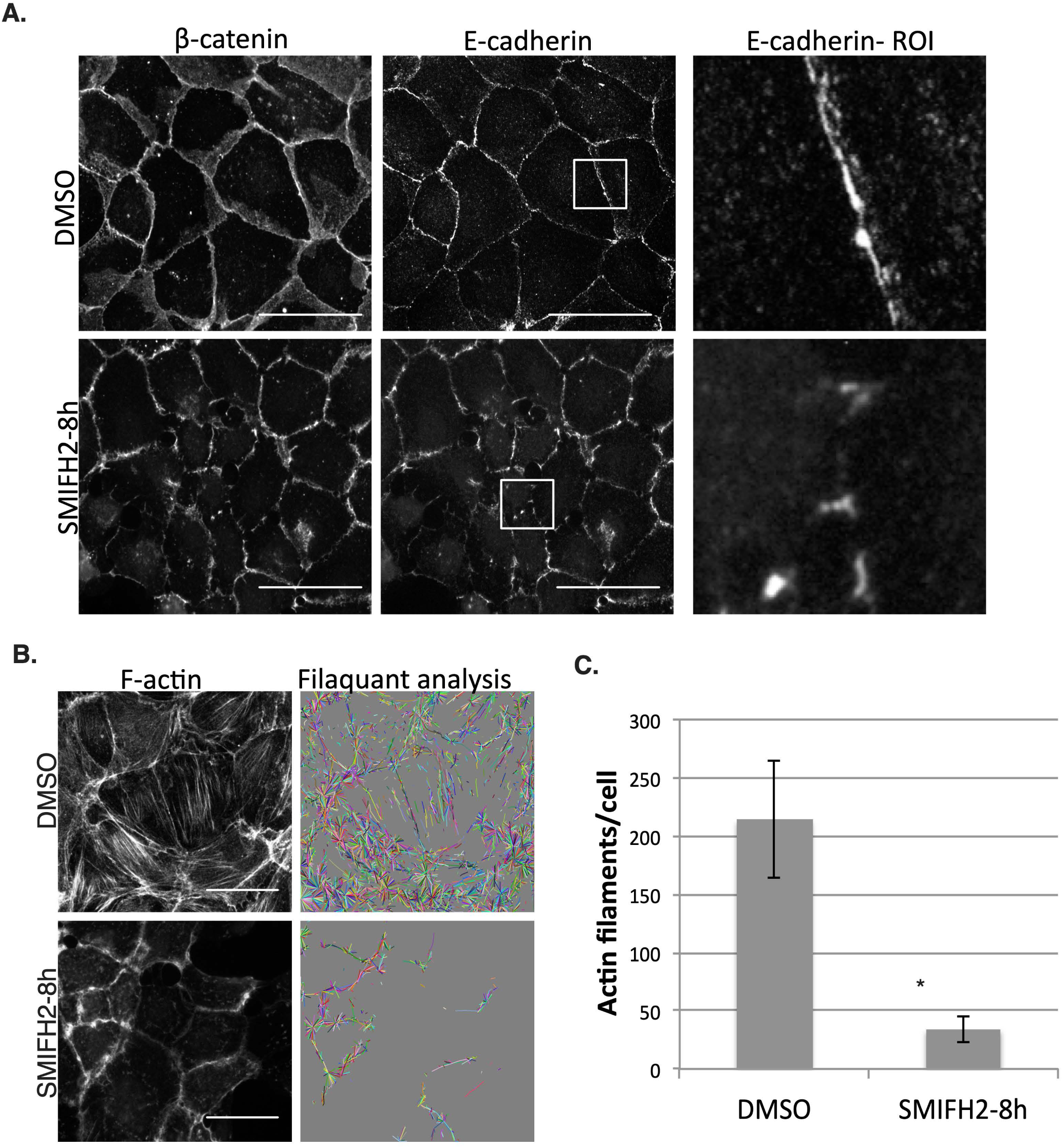
Formin inhibition disrupts AJs and decreases F-actin filaments. OVCA429 cells were treated with 40 μM SMIFH2 or DMSO for 8 hours. **A**. Junctions are stained for β-catenin and E-cadherin. Rectangles mark regions of interest (ROIs). Scale bars = 50 μm. **B**. F-actin was stained with phalloidin and detected filaments shown in Filaquant software analysis. **C**. A significant reduction in F-actin is observed in SMIFH2-treated cells. Five random fields and at least 58 cells were analyzed per condition. *p<0.05. Values denote mean ± SEM.

